# Phylogenetic analysis of pathogenic and non-pathogenic *Trichoderma* isolates from plants, soil and commercial bio products

**DOI:** 10.1101/2024.09.27.615377

**Authors:** Annette Pfordt, Clovis Douanla-Meli, Bernhard C. Schäfer, Gritta Schrader, Eike Tannen, Madhav Jatin Chandarana, Andreas von Tiedemann

**Affiliations:** Plant Pathology and Crop Protection, Georg-August-University of Goettingen, Goettingen, Germany; Julius Kühn-Institut (JKI) - Federal Research Centre for Cultivated Plants, Institute for National and International Plant Health, Germany

**Keywords:** Trichoderma, biocontrol, maize ear rot, *T. afroharzianum*, *T. harzianum*, *TEF1-α*

## Abstract

Fungi of the genus *Trichoderma* are found worldwide in various types of soil, plant rhizospheres, and plant materials. Several *Trichoderma* spp. are used in crop health management to promote growth and control plant diseases. Although widely considered beneficial, some members have been reported to be pathogenic to maize, causing a disease called Trichoderma ear rot. Since 2018, *T. afroharzianum* has caused significant infections of maize cobs in Germany, France and Italy. This study aimed to investigate the pathogenicity and phylogenetic relationships among different *Trichoderma* strains from diverse sources and geographical origins. Species identification and phylogenetic analysis were performed by sequencing *internal transcribed spacer* (*ITS*), *translation elongation factor 1-α* (*TEF1-α*) and *RNA polymerase II subunit B* (*RBP2*) genes, and pathogenicity was tested by artificially inoculating maize cobs under controlled greenhouse conditions. A total of 131 isolates were analyzed and assigned to 20 *Trichoderma* species. Among these, 39 isolates from six species were pathogenic, causing symptoms of green spore layers between kernels and husk leaves. While previous studies primarily identified *T. afroharzianum* as the main species causing Trichoderma ear rot, this study found that isolates of *T. asperellum*, *T. atroviride* and *T. guizhouense* also exhibit pathogenicity on maize cobs. Additionally, *Trichoderma* strains from commercial biocontrol products displayed unexpected pathogenicity inducing up to 92% disease severity on maize cobs. Most *T. afroharzianum* strains induced high levels of disease severity, although some isolates of the same species did not cause any disease, indicating a large heterogeneity in pathogenicity within the species. Notably, phylogeny reconstruction based on the *TEF1-α* and *RBP2* genes, did not result in any discernible clustering between pathogenic and non-pathogenic isolates. A further novel finding is the isolation of pathogenic *Trichoderma* isolates from soil, demonstrating that soil can serve as a reservoir for pathogenic species. This study highlights the need for careful selection and monitoring of *Trichoderma* strains for agricultural use, considering their beneficial and pathogenic potential.

**Author Summary:** In this study, we explored the ability of different *Trichoderma* species to infect maize plants. *Trichoderma* is a group of fungi known for its beneficial role in agriculture, often used as a biological pesticide to control fungal plant diseases. However, some species within this group can also act as pathogens, causing infections in crops like maize. We found that one species, *T. afroharzianum*, is particularly aggressive, capable of infecting maize without the plant being wounded first. This makes it a potentially serious threat to crop health. In contrast, other species, such as *T. atroviride* and *T. asperellum*, only caused infections when the maize was already damaged. Our research suggests that pathogenic *Trichoderma* species not only effectively infect plants but can also survive well in soil, making their control difficult. These findings highlight the need for better understanding of how these fungi operate in order to manage the risks they pose to important crops like maize, while still taking advantage of their beneficial uses in agriculture.

## Introduction

Members of the genus *Trichoderma* (Ascomycota) are opportunistic fungi capable of rapidly colonizing diverse niches in both natural and artificial environments. In these habitats, *Trichoderma* spp. contribute to nutrient cycling by decomposing complex organic compounds, thus enhancing soil fertility and promoting overall ecosystem health (1,2). Beyond their ecological functions, *Trichoderma* spp. are used in agriculture as biocontrol agents to promote plant health by directly controlling phytopathogens or indirectly enhancing plant growth (3–5). *Trichoderma*-based products are used to manage soil and foliar diseases on maize and other important crops. Degani and Dor (2021) determined a reduction of plant mortality and improvement of growth parameters of maize seedlings against *Magnaporthiopsis maydis* after the application of *T. asperelloides* and *T. longibrachiatum* in the soil (6). Several studies reported a positive effect against *Fusarium* infection in maize cobs and stalks after treating seedlings with *Trichoderma* spore suspension or applying seed coatings to reduce damping-off caused by *Pythium* and *Fusarium* species (7).

However, though widely considered beneficial, members of the *Trichoderma* genus have been also reported to be pathogenic causing damage and yield losses to maize (8–10). The earliest report of a *Trichoderma* species pathogenic on maize has been released from the University of Kansas in 1910, where *T. viride* was described as a greenish-yellow wet mold growing between the rows of maize kernels (10). The infection frequently resulted in the germination of grains on the cob and reportedly occurred widely on some fields in Riley County (Kansas, USA). *Trichoderma* species were also identified to act as secondary pathogens, infecting the root and stalk of maize after preceding infection by *Fusarium oxysporum* in Ontario, Canada (11). In 1972, *T. koningii* was isolated from stunted maize plants in southern Ontario (12) and three years later, McFadden and Sutton (1975) *T. koningii*, *T. harzianum*, and *T. hamatum* were reported to be the causal agents of lesions on the first internode of maize (13).

The first mention of *Trichoderma* associated with the maize ear rot disease was published by the University of Illinois in 1991 (14). Thereby, *T. viride* was described as the causal agent of maize ear infection after insect or mechanical damage. Similar results were published as farm management reports, several years later in Iowa (15), Ohio (16) and Kentucky (17), describing *Trichoderma* as an ear rot disease in maize with dark, gray-green conidial layers between the kernels of infected maize cobs after insect feeding or mechanical damage. Infections occasionally resulted in massive outbreaks with premature germination of kernels (15,16). In additions to maize, *T. afroharzianum* has been reported to cause disease symptoms and yield losses in other cereals such as wheat and barley (18).

In 2020, the disease was reported for the first time in Europe (8). In this report, *T. afroharzianum* was identified as the causal organism of maize ear rot infection in Southern Germany. *T. afroharzianum* is one of the morphologically similar species phylogenetically clustering in the *Harzianum* Clade of *Trichoderma*. *T. afroharzianum* is widely distributed and commonly found as an endophyte (19). Trichoderma infection in maize was characterized by mycelial growth with colored green spores in the inter-kernel regions and on the outer surface of the husks. However, the disease was not associated with bird feeding or mechanical injuries. After inoculation in the field and greenhouse, a significant reduction in the dry weight of maize cobs and premature germination of kernels was reported (8). Cob weight was significantly reduced by up to 50% compared to control plants due to the enzymatic activity of *T. afroharzianum*, particularly the production of alpha-amylase, which resulted in the degradation of starch into simpler sugars such as glucose. This lead to negative effects on seedling development by causing premature germination, reduced germination rates, and implications for agricultural food and feed production (20).

Recently, Trichoderma ear rot caused by *T. afroharzianum* has also been reported from France and Italy (9). Pfordt et al (2020) showed differences in pathogenicity within the *T. afroharzianum* species, with reference to type strain CBS124620 which caused no symptoms after artificial inoculation while other *T. afroharzianum* species showed high pathogenicity on maize cobs. This indicated phylogenetic separation on the species level into pathogenic and non-pathogenic strains and suggests that further research is required to clarify the genetic differences between these strains within the species *T. afroharzianum* (8).

Therefore, the aim of the study was to identify pathogenicity within the *Trichoderma* genus and relate pathogenicity to the phylogenetic position of different *Trichoderma* species and strains originating from diverse sources and origins, such as plants, soil, and commercial biocontrol products. A specific aim of the study was to determine whether other species of *Trichoderma* besides *T. afroharzianum* can be pathogenic on maize. The phylogenetic differentiation was based on sequencing the *translation elongation factor 1-α* (*TEF1*-α), *internal transcribed spacer* (*ITS*) and *RNA polymerase II subunit B* (*RBP2*) genes. A further objective was to assess whether *Trichoderma*-based biocontrol products can induce disease symptoms on maize cobs. Finally, the study aimed to determine whether pathogenic species can be isolated from soil, emphasizing the role of soil as a reservoir for inoculum of these pathogens.

## Results

### Collecting fungal isolates and identification

A total of 153 distinct strains were successfully isolated from various sources and geographical locations. Additionally, 60 isolates were obtained from the soil of maize fields, while eight isolates were collected from the rhizosphere of various plants. Furthermore, 18 isolates derived from various plant material and cultivated mushrooms. Geographically, the majority of isolates was collected in Germany (69 isolates), but further isolates were obtained from diverse origins like Serbia (25 isolates), France (11 isolates), Macedonia (2 isolates), China (2 isolates), and Italy (1 isolate). In order to verify the identity of the isolates from the different sources as *Trichoderma* spp., single spore cultures were carefully prepared and their morphological characteristics were subjected to a comparative analysis. Figure 2 shows single spore colonies obtained from 16 Trichoderma isolates after eight days of cultivation on PDA. The colony color ranged from green to dark green, yellow to greenish and white to pale yellow. The intensities of yellow and green were different among these isolates. Isolates of species in the *Harzianum* clade, and especially *T. afroharzianum, T. guizhouense*, produced diffusible yellow and light green pigments. Isolates belonging to the *Viride* clade, *T. atroviride, T. asperellum, T. koningii*, produced grayer and darker green pigments. Pigmentation of *T. atrobrunneum* isolates was lighter in comparison to other species with white to pale yellow colors (Figure 1).

**Figure 1:**
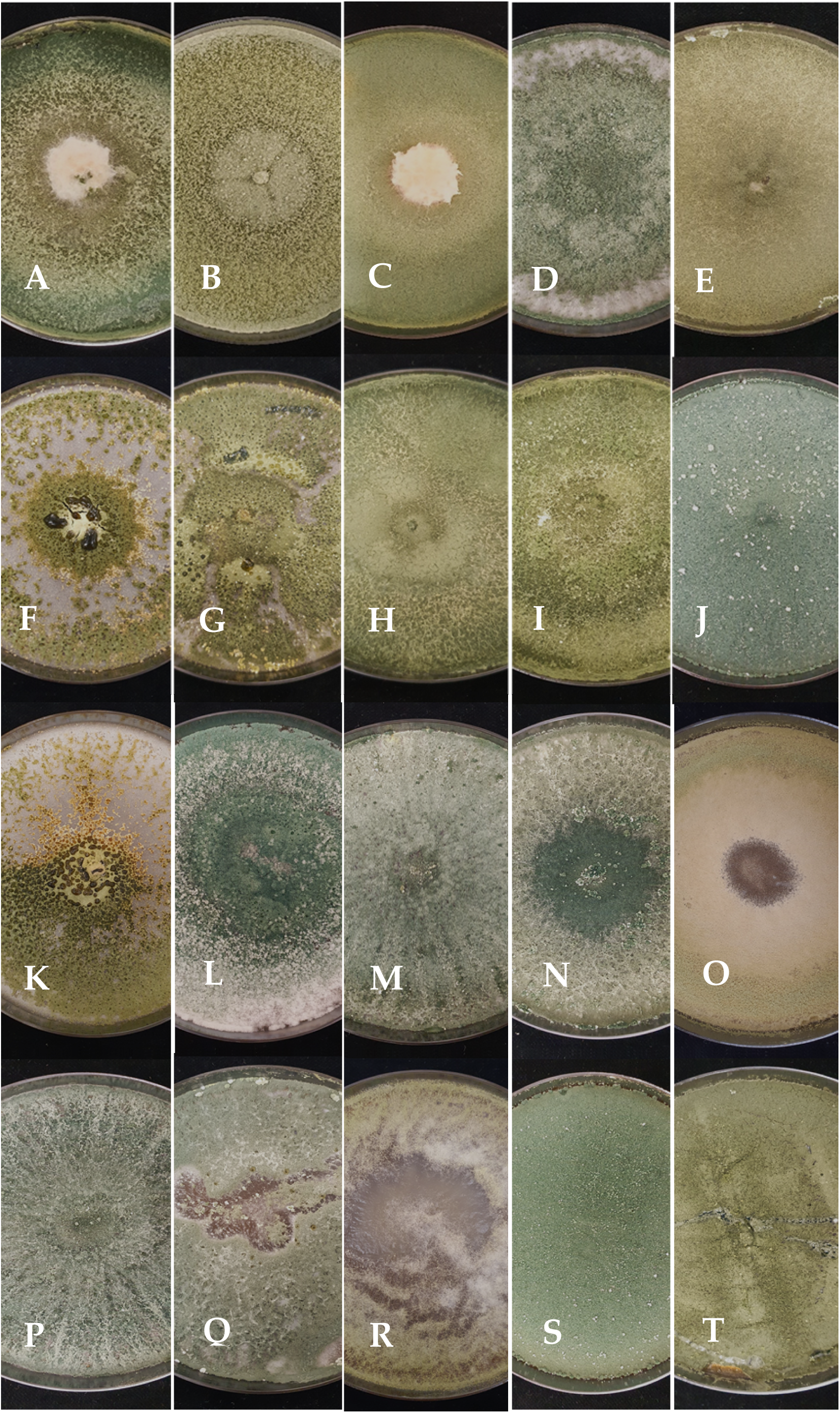
Colony characteristics of various *Trichoderma* species on PDA plates, after ten days after growth at 22°C in growth cambers (A) AP18TRI2, *T. afroharzianum;* (B) AP18TRI4, *T. tomentosum*; (C) AP19TRI5, *T. afroharzianum;* (D) IPP0316, *T. atroviride,* (E) IPP0319, *T. harzianum*; (F) AP19TRI12, *T.harzianum;* (G) IPP1663, *T.harzianum*, (H) IPP1657, *T. koningii*; (I) CBS124620, *T. afroharzianum*, (J) TR4, *T. asperellum*; (K) TR2, *T. guizhouense*; (L) TR10, *T. atroviride*, (M) VINTEC_SC1, *T. atroviride,* (N) MRI349, *T. afroharzianum*; (O) TR9 *T. atrobrunneum*; (P) T60, *T. atroviride;* (Q) TRISOIL_I1237*, T. atroviride;* (R) TR9, *T.atrobrunneum*; (S) TR4, *T. asperellum;* (T) XILONT34, *T. asperellum*.

**Figure 2.**
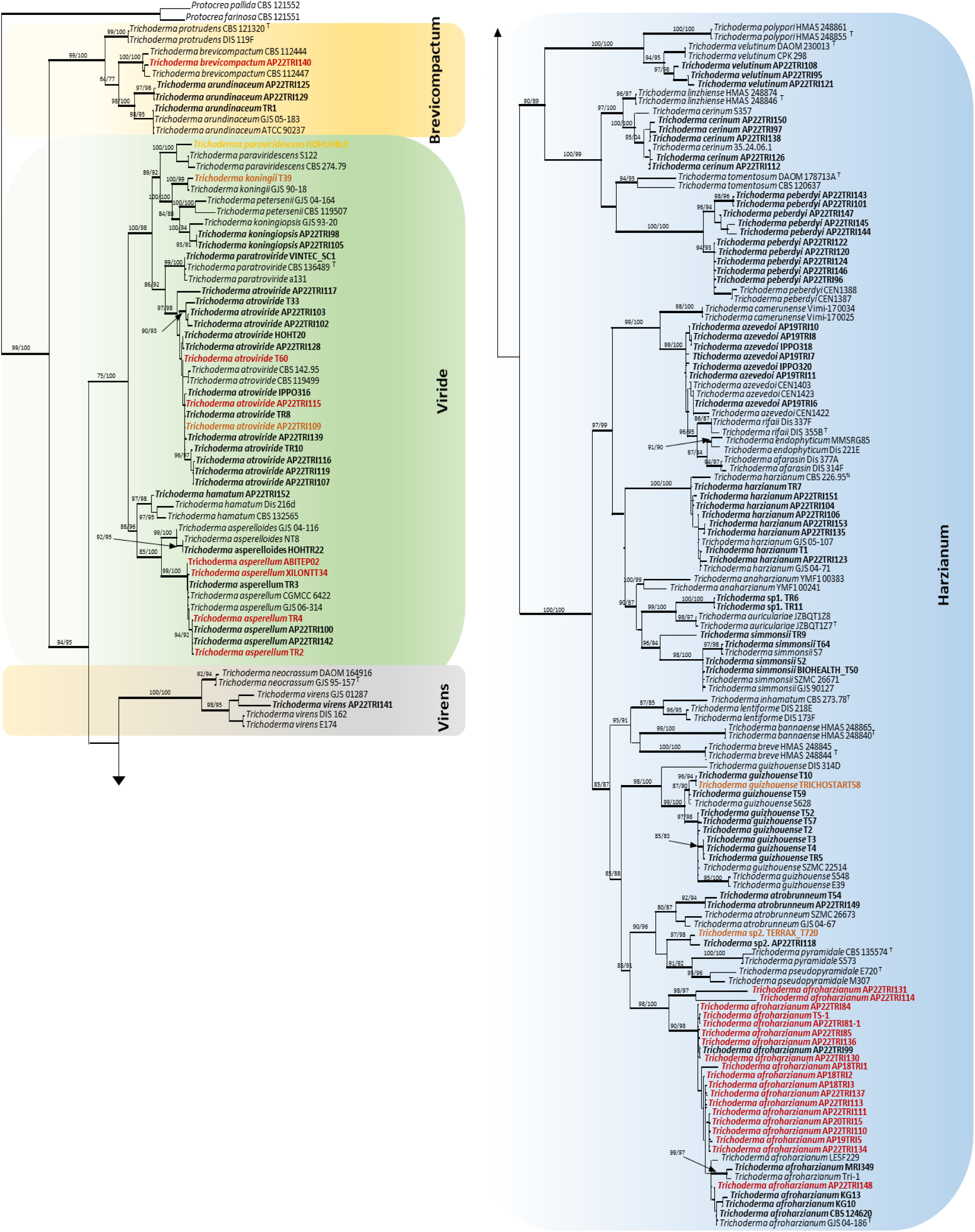
Phylogenetic tree based on Maximum Likelihood analysis of a concatenated *TEF1-α* and *RPB2* sequence dataset. Bootstrap values higher than 70 % from MP (left) and RAxML (right) analyses are given above the nodes. Thickened lines represent branches with a Bayesian posterior probability greater than 0.95. Isolates analyzed and tested for pathogenicity on maize in this study are in bold. Pathogenic isolates are in bold color with red for high, brown for moderate and orange for weak disease severity. *Protocrea farinosa* (CBS 121551) and *Protocrea pallida* (CBS 121552) were used as outgroup.

### Phylogenetic analyses and taxonomy

Preliminary BLAST searches with *ITS* sequences identified 22 isolates from maize plants and 62 from maize field soil as belonging to *Trichoderma*. Including *Trichoderma* from other sources, *TEF1-α* and *RPB2* sequences of a total of 113 *Trichoderma* isolates were determined for the phylogenetic analyses. The phylogenetic taxonomy delimited 20 distinct *Trichoderma* species. Among the isolated strains, *T. afroharzianum* was most frequently isolated from infected maize plants and soil with 29 distinct isolates. Similarly, *T. asperellum* (7 isolates), *T. atroviride* (18 isolates), and *T. guizhouense* (10 isolates) were also notably present. The isolation of four strains of *T. atrobrunneum*, three strains of *T. simmonsii*, and three strains of *T. hamatum* further contributed to the multifaceted composition of the isolated strains. Additional species within the range included *T. tomentosum* (5 isolates), *T. brevicompactum* (3 isolates)*, T. peberdyi* (10 isolates), and unique representatives of *T. koningii, T. asperelloides, T. paraviridescens, T. arundinaceum, T. velutinum, T. koningiopsis, and T. virens* (one isolate each).

The combined data of *TEF1-α* and *RPB2* consisted of 199 entries with *Protocrea farinosa* (CBS 121551) and *Protocrea pallida* (CBS 121552) used as outgroup. These concatenated data comprised 2,520 characters including gaps, with 1,406 characters for *TEF1-α* and 1,114 characters for *RPB2*, among these, 880 (34.4 %) characters were parsimony-informative for MP analyses. All concatenated trees constructed in MP, ML and BI analyses presented nearly similar topologies and the ML tree was selected and shown in Figure 2. The individual *TEF1* and *RPB2* trees (Supplementary Figures A1, A2) well separated *Trichoderma* species providing almost the same groups resolved by the concatenated trees. For specific resolution, RPB2 tree was highly congruent with concatenated trees. However, resolution of some species differed in the TEF1-α tree, such as T. tomentum with the two isolates placed, but separated, to *T. peberdyi* and *T. azevedoi* with its isolates separated into two distant groups (Supplementary Figure A1). All concatenated trees constructed in MP, ML and BI analyses presented nearly similar topologies and the ML tree was selected and shown in Figure 2. Phylogenetic analyses grouped all *Trichoderma* sequences into four known clades, namely the *Brevicompactum* clade, the *Harzianum* clade, the *Virens* clade and the *Viride* clade. Among *Trichoderma* isolates, 77 isolates were placed in the species-rich *Harzianum* clade and assigned to *T. afroharzianum*, *T. atrobrunneum*, *T azevedoi*, *T. cerinum*, *T. guizhouense*, *T. harzianum*, *T. peberdyi*, *T. simmonsii* and *T. velutinum*. Additionally, four isolates (TERRAX_T720, AP22TRI118, TR6 and TR11) in the *Harzianum* clade were resolved in two distinct branches of two isolates each, showing close affinity to *T. pyramydale* and *T. auriculariae,* respectively. Thirty-one isolates were resolved in the *Viride* clade and belonged to *T*. *atroviride, T. hamatum, T. koningii, T. koningiopsis, T. paratroviride, T. paraviridescens, T. asperelloides* and *T. asperellum*. One species clustered as *T. virens* in the *Viride* clade and two species were identified as *T. arundinaceum* and *T. brevicompactum* in the *Brevicompactum* clade.

*Trichoderma* isolates associated with maize in the field grouped in the *Harzianum* clade and belonged to species like *T. afroharzianum* (isolates AP18TRI1 AP18TRI2, AP18TRI3, AP18TRI5, AP19TRI15, AP19TRI16, AP19TRI17, AP22TRI81_1, AP22TRI19, AP22TRI20), *T. azevedoi* (isolates AP19TRI6, AP19TRI7, AP19TRI8, AP19TRI9, AP19TRI10, AP19TRI11, AP19TRI12) and *T. harzianum* (isolate AP19TRI14, AP19TRI9, AP19TRI12). In general, isolates showing pathogenicity on maize during greenhouse trials scattered in the *Harzianum* and the *Viride* clades (Figure 3). Pathogenicity was not consistently corroborated for all tested isolates of any *Trichoderma* species and all phylogenetic analyses failed to distinguish between pathogenic and apathogenic isolates of *T. afroharzianum*, *T. asperellum, T*. *atroviride* and *T. guizhouense*.

**Figure 3:**
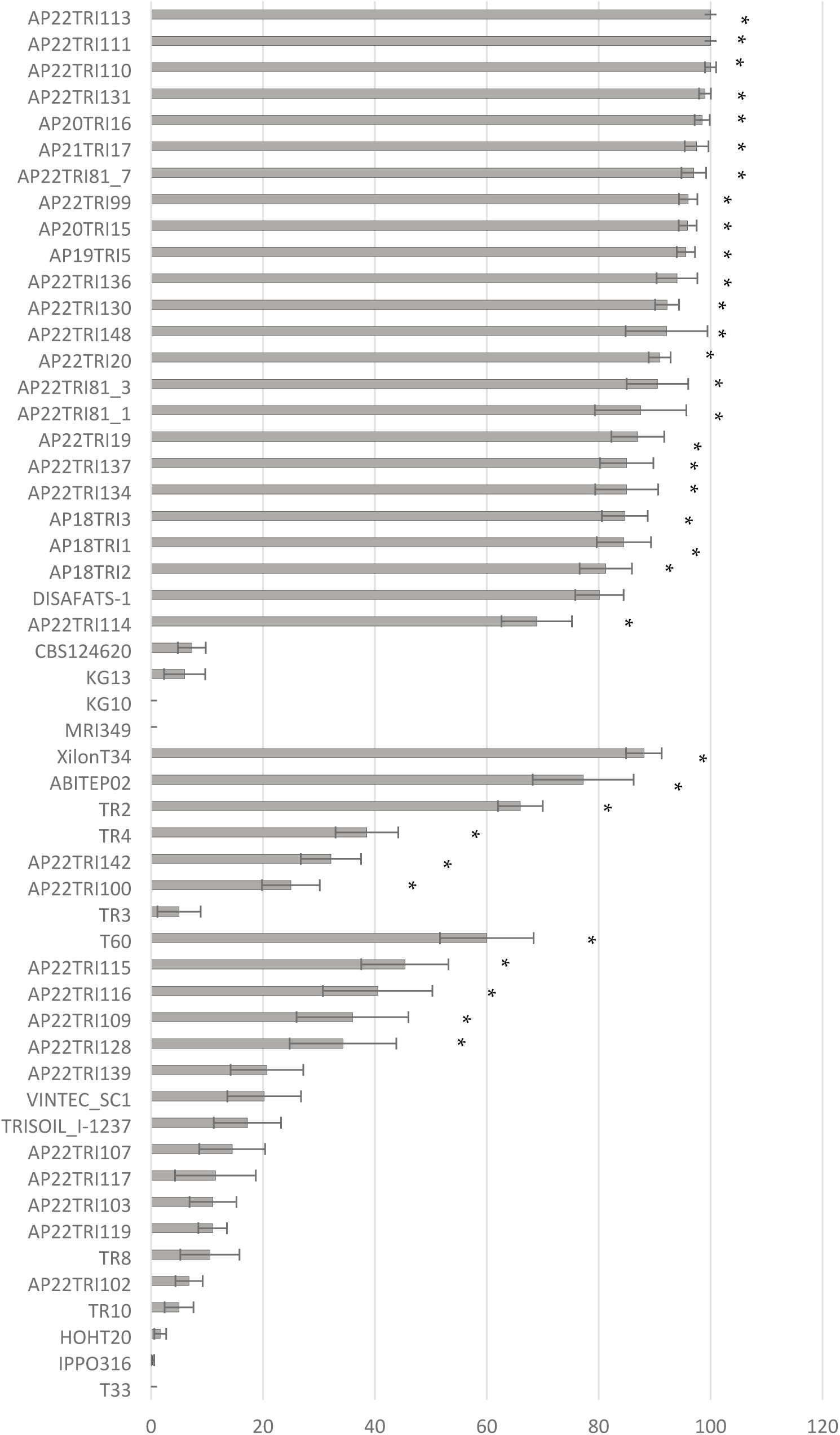
Mean disease severity (%) of isolates of *T. afroharzianum*, *T. asperellum* and *T. atroviride* on maize cobs, 28 days after artificial inoculation in the greenhouse. Bars represent standard error. Asterisks (*) indicate significant differences to water inoculated control plants (Tukey-Test, p ≤ 0.05).

### Disease severity

Four weeks after inoculation, typical symptoms of Trichoderma ear rot were observed and disease severity was assessed visually. A total of 133 isolates from 23 *Trichoderma* species were screened for their pathogenicity on maize cobs in the greenhouse. Twenty-nine *T. afroharzianum* isolates were tested during greenhouse pathogenicity tests with a mean disease severity of 78%. Of those, 25 isolates caused disease severities of 70 to 100%, while the isolates CBS124620 (type strain of *T. afroharzianum*), AP22TRI121 and KG10 caused only light infection (< 10%), not significantly different to the water control. No infection was observed from isolates MRI349 and KG10.

A total of seven *T. asperellum* isolates were tested in the greenhouse, causing an average disease severity of 47.4%. Among the evaluated isolates, three isolates of *T. asperellum* caused moderate to high infection rates ranging from 66% to 88% disease severity on maize cobs. T34, which serves as the active component in the biofungicide Xilon® (Kwizda Agro GmbH), caused the highest disease severity, resulting in a disease severity level of 88%. Isolate ABITEP02, an approved soil additive from the company AbiTep®, caused 77% infection, followed by TR2 (66%). Isolate TR4, isolated from apricot in Serbia, and isolates APP22TRI142 and APP22TRI100, isolated from soil, caused moderate infections with 25-38% disease severity.

Eighteen *T. atroviride* isolates were isolated, causing an average disease severity of 19.9%. Five isolates caused significantly higher disease severity than water-inoculated controls, with mean infections ranging from 34% to 60%. Twelve isolates caused light infections of <20% with no significant differences from water-inoculated controls. A total of 8 *T. harzianum* isolates were tested for pathogenicity in the greenhouse, however, none of the isolates led to significantly higher disease severity than water inoculated control plants. Disease severity ranged from 0 to 14% with an average severity of 2.3%. Two isolates of *T. atrobrunneum* were tested, however, only one isolate induced a notable infection level of 27%. Ten isolates of *T. guizhouense* were tested, resulting in an average disease severity of 4.8%. Only one isolate showed significant infection with an average disease severity of 42%. The remaining nine isolates did not exhibit any significant infection. Three isolates of *T. simonsii*, one isolate of *T. hamatum*, five isolates of *T. cerinum*, seven isolates of *T. azevedoi* and ten isolates of *T. peberdyi* caused no or only mild symptoms, not significantly different from the water control. Among the isolates of *T. koningii, T. asperelloides, T. paraviridescens, T. arundinaceum, T. velutinum, T. koningiopsis,T. virens,* and *T. brevicompactum* tested in the greenhouse, only one isolate of *T. brevicompactum* (AP22TRI140) induced a significant level of disease severity of 37% (Figure 4).

**Figure 4:**
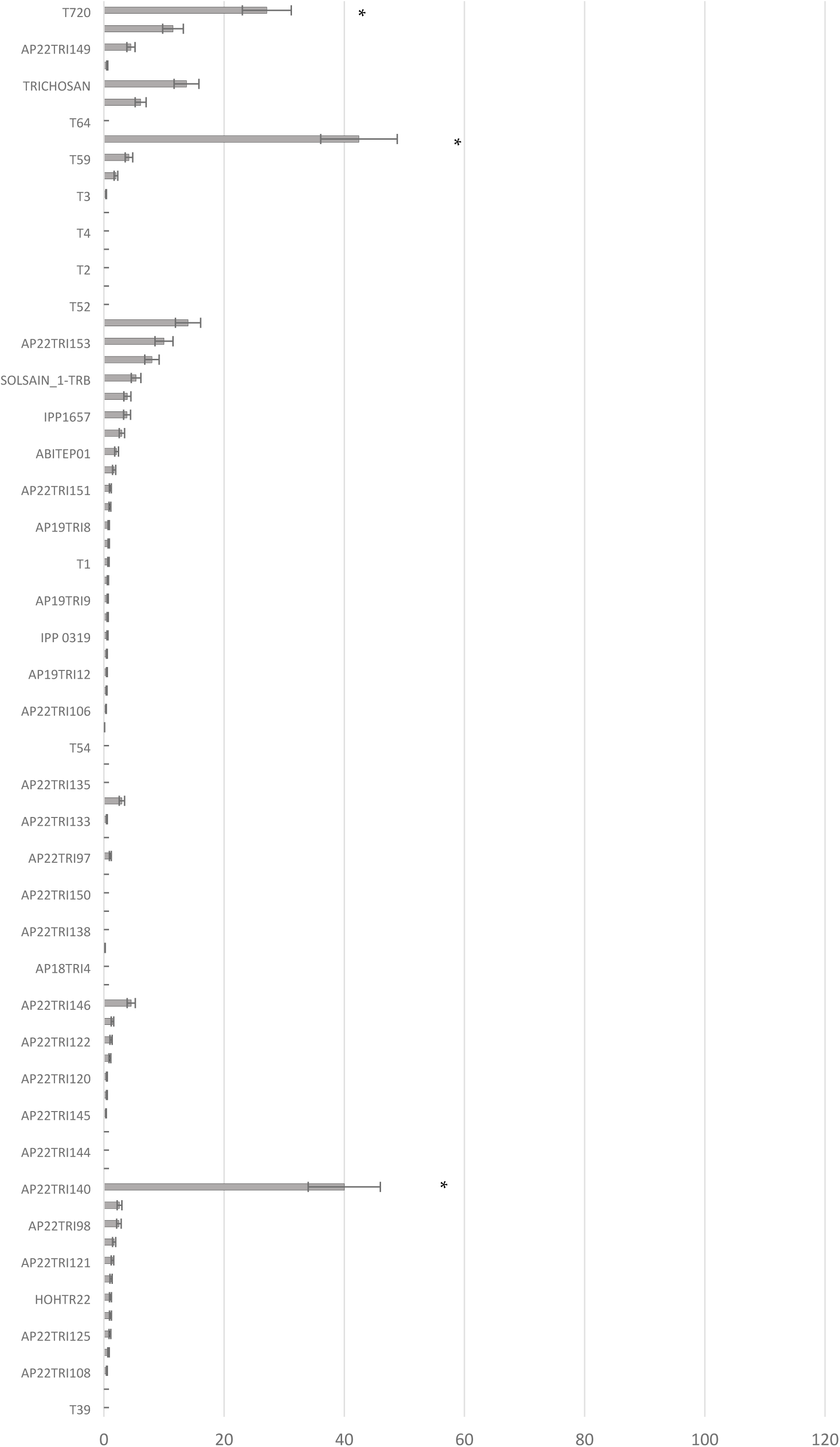
Mean disease severity (%) of collected isolates of *T. atrobrunneum*, *T. simonsii*, *T. guizhouense*, *T. harzianum*, *T. hamatum*, *T. linzhiense*, *T. tomentosum*, *T. peberdyi* and various species on maize cobs, 28 days after artificial inoculation in the greenhouse. Bars represent standard error. Asterisks (*) indicate significant differences to water inoculated control plants (Tukey-Test, p ≤ 0.05).

The highest diversity of *Trichoderma* species was isolated from soil. In total, 13 different species were identified, with the most common being *T. afroharzianum* (12 isolates), *T. atroviride* and *T. peberdyi* (10 isolates each), followed by *T. harzianum* (7 isolates). Notably, most *T. afroharzianum* isolates were obtained from soils in locations with previous observations of Trichoderma ear rot infections, particularly in southern Germany, such as Haßloch, Rustenhart (France), and Oberpframmern. Many of these isolates displayed high pathogenicity (>70% disease severity) on maize cobs after artificial inoculation in the greenhouse. However, one isolate from Nossen (AP22TRI121) exhibited no pathogenicity. Interestingly, pathogenic isolates of *T. afroharzianum* were also found in northern locations without any prior reports of Trichoderma ear rot, such as in Bevern (AP22TRI99). *T. atroviride* and *T. asperellum* isolates from soil displayed medium to low pathogenicity, while none of the other species isolated from soil were pathogenic on maize cobs.

The isolates were collected from 48 different locations and origins. The majority of pathogenic isolates originated from plants and soil in Southern Germany and France, as well as from biological plant protection products. In contrast, most apathogenic isolates were collected from plants in Serbia and Northern Germany.

Among the ten strains from approved biocontrol and biostimulant products, strain T34 from the product Xilon® (Kwizda Agro GmbH) exhibited the highest pathogenicity (88% disease severity), followed by AbiTep02 (AbiTep) (77.2%) and strain T58 from Trichostar® (Intrachem Bio Deutschland GmbH & Co) (42.8%). Trichoderma strains from other bio-products showed only mild or no pathogenicity, comparable to the water control plants.

## Discussion

In this study, 153 isolates of *Trichoderma* spp. were obtained from maize plants, soil of maize fields and commercial biocontrol products. The identity of the isolates was determined using sequences of *ITS*, *TEF1-α*, and *RPB2* genes and their ability to cause ear rot on maize was tested under controlled conditions in the greenhouse.

Previous reports focused on *T. afroharzianum* as the main causal species of Trichoderma ear rot in maize (8,9). While *T. afroharzianum* was the predominant species isolated from naturally infected maize cobs, our present investigation demonstrated that other *Trichoderma* species obtained from other plant materials and soil may also exhibit pathogenicity after needle pin inoculation of maize cobs. This demonstrated significant differences in the pathogenic potential and infection mechanisms of various Trichoderma species on maize cobs. Notably, *T. afroharzianum* is capable to infect maize tissues without prior wounding, indicating its advanced ability to overcome the plant physical barriers and immune defenses (21,22). In contrast, strains of *T. atroviride* and *T. asperellum* were found to require wounds or entry points such as after insect feeding or mechanical injury to initiate infection. This distinction emphasizes the importance of understanding the diversity of host-pathogen interactions exhibited by Trichoderma, in particular the transition from wound-dependent to fully competent pathogenicity.

The majority of *T. afroharzianum* strains exhibited high disease severity of 75% or above. However, other isolates of the same species, such as KG10, KG13, MRI349 and AP22TRI99 did not cause any disease on maize cobs. This further indicates a significant heterogeneity among *T. afroharzianum* strains, varying between pathogenic and non-pathogenic life styles. One explanation for this phenomenon could be differential expression of pathogenicity factors among these isolates. Pathogenicity in fungi often involves the production and secretion of various enzymes and metabolites that facilitate host invasion and colonization (23,24). It is conceivable that pathogenic isolates of *T. afroharzianum* possess genetic determinants that enable them to express these factors at higher levels or under specific environmental conditions, leading to pathogenicity on maize cobs (25). Conversely, non-pathogenic isolates may lack or exhibit reduced expression of these pathogenicity factors, resulting in inability to cause disease (26). Reports on *Trichoderma* as a plant pathogen are relatively seldom compared to its beneficial interactions with plants. The factors that allow some *Trichoderma* strains to infect plants are poorly studied. This underscores the complexity of Trichoderma-host interactions and suggests that genetic variability within *T. afroharzianum* populations may determine their pathogenic potential.

However, we were unable to differentiate between pathogenic and nonpathogenic isolates within the *T. afroharzianum* species using phylogenetic analyses based on *TEF-α* and *RBP2* genes. The reconstructed phylogeny with the combined *TEF-α* and *RBP2* was fully resolved with strongly supported branches. However, *ITS* was quite inaccurate and did not contribute to species identification of *Trichoderma.* While *ITS* is only useful for differentiation on the genus level (27), *TEF-α* and *RBP2* are well conserved and have a much higher phylogenetical power to discern *Trichoderma* species (19,28,29). Nonetheless, using sequences of the two genes, it was not possible to distinguish between pathogenic and non-pathogenic isolates of *T. afroharzianum*, *T. asperellum*, *T. atroviride* and *T. guizhouense*. Hence, it seems to be very challenging to predict pathogenicity from such sequence polymorphisms because pathogenicity factors are typically controlled by a complex interplay of multiple genes, including those involved in the production of enzymes, secondary metabolites, and other virulence factors (30,31). Presumably, the genetic determinants of *Trichoderma* pathogenicity are likely located in genomic regions outside the *TEF-α* and *RBP2* genes, such as accessory genomes or specific pathogenicity islands, which were not part of this analysis. Therefore, a stronger focus on the genomic level will be important to understand pathogen-host interactions and thereby elucidate the origin of pathogenicity of *Trichoderma* causing maize ear rot. Some previous studies revealed that effectors may play a major role in pathogenicity by overcoming plant immunity and establishing the interaction with the host (32,33). In the same vein, carbohydrate active enzymes (CAZymes) which are important for pathogens to retrieve carbohydrates from host tissues, can determine the fungal lifestyle (34–36). Both, effectors and CAZymes, may be shared in the same profile by isolates or species colonizing the same host. This indicates that genomic differences could explain the differences between species or isolates that exhibit different lifestyles (37).

Aggressiveness of can also be influenced by environmental factors such as temperature, humidity, and precipitation. Climatic conditions can directly influence growth, development and occurrence of *Trichoderma* spp. in the environment (38,39). Most pathogenic isolates were isolated from locations characterized by warm temperatures and dry conditions like Southern Germany, France and Italy. It is possible that pathogenic isolates of *T. afroharzianum* have adapted to specific environmental conditions, allowing them to infect maize cobs effectively. Further research using whole-genome sequencing, transcriptomics, or proteomics will be necessary to identify the specific genetic or regulatory factors that differentiate pathogenic and nonpathogenic isolates within the *T. afroharzianum* species.

In this study, the pathogenicity of *Trichoderma* strains contained in three biological plant protection products and six biostimulants was evaluated on maize cobs. Three strains were pathogenic and induced medium to high disease severity of up to 85% after inoculation in the greenhouse, while the other strains were non-pathogenic on maize.

*Trichoderma* spp. are widely used as actives in biocontrol products to manage plant pathogens, as they exhibit antagonistic properties against a broad range of soil-borne and foliar pathogens, such as *Fusarium*, *Rhizoctonia*, *Pythium*, and *Sclerotinia* (40,41). Biocontrol products primarily aim to protect crops from pests and diseases to preserve crop health and are considered as a potential alternative to chemical pesticides. The registration of biological fungicides involves a risk assessment to exclude unacceptable risks to human health, non-target organisms, the crop plant or the environment. It is noteworthy, that there are several reports of negative effects of biocontrol agents on non-target organisms and the environments (42–44). Biocontrol agents are in general selected for traits that enhance their survival, competitiveness, and efficacy against pests (45). After application, this divergent selection pressure can inadvertently favor strains that possess pathogenic traits, particularly if those traits provide a survival advantage in certain conditions (42). This evolutionary shift and chance of nutritional habits from a mycoparasite to a plant mutant over the course of evolution has already been described. This genus has constantly redesigned its genome to improve its ability to rapidly colonize and successfully compete in novel habitats (46). It is only natural that *Trichoderma* will adapted and find new nutritional source by infecting plants as a pathogen.

Genetic mutations after application in soil can occur naturally within populations of biocontrol agents which might enhance pathogenicity by enabling the agent to better infect and colonize host plants (47). These mutations can arise spontaneously and, if beneficial for survival or reproduction, may become more prevalent in the population (48). In addition, fungi which are used as biocontrol agents can adapt to specific ecological niches and develop traits that allow them to exploit new resources or hosts (49).

However, it is important to mention that the pathogenic *Trichoderma* strains from biocontrol products used in this study, were tested only after artificial inoculation in the greenhouse, where spores were directly applied to maize cobs. It remains unknown whether these pathogenic strains from bio-products could infect maize cobs also under natural conditions or whether they can spread to maize cobs after soil application. Further studies are needed to determine their behavior and the impact under agricultural settings. In addition, the identification of pathogenic strains among commercial products highlights the need for rigorous quality control measures in the production and application of these products. Biostimulants are registered under the Fertilizing Products Regulation (EU) 2019/1009 which does not require any proof of efficacy and has rather low level safety requirements (50).

A further novel finding from this study is the successful isolation of pathogenic *Trichoderma* strains from soil samples, demonstrating soil to be a reservoir for *Trichoderma* inoculum. A total of 60 soil samples from 19 locations with and without previous observation of Trichoderma ear rot infection were analyzed. Pathogenic *T. afroharzianum* were recovered from the soil at all sites where Trichoderma ear rot had previously occurred. This suggests that the soil is a potential source of inoculum for cob infection. However, it is notable that pathogenic isolates were also detected in some areas without a cob infection history, indicating that *Trichoderma* is present in the soil but maybe lacks the necessary conditions for infection.

*Trichoderma* species exist worldwide in various environments as saprophytes in the soil obtaining nutrients from dead or decaying organic matter (51,52). *Trichoderma* species can survive for varying lengths of time depending on environmental conditions, competition with other microorganisms, and the availability of organic matter. Generally, they can persist in soil for several weeks to months, especially if conditions are favorable for their growth (53,54). Pathogenic *Trichoderma* strains are capable of surviving in the soil by efficiently colonizing organic material, similar to saprophytic strains. This saprophytic competence allows them to remain viable in the absence of a host plant, utilizing decaying organic matter as a nutrient source. Consequently, even when crop plants are not present, these pathogenic strains can survive and multiply in the soil, ensuring their persistence over time. This dual ability to act as both pathogens and saprophytes provides these strains with superior advantages to establish in a field ecosystem to induce continuous or recurrent infections. *Trichoderma* species lack sexual reproduction, therefore asexual sporulation by conidia is a common process of reproduction (55). In order to survive and spread, *Trichoderma* switches from vegetative to reproductive development induced by light and mechanical injury (56). Pathogenic *Trichoderma* strains can reproduce effectively through conidia formation, similar to saprophytic strains. This reproductive strategy ensures their widespread dissemination and survival in various environmental conditions. The combination of efficient reproduction and pathogenicity makes these strains particularly dangerous, as they can spread rapidly and establish infections under favorable conditions.

It is still unknown how pathogenic *Trichoderma* species infect maize cobs in the field but it can be assumed that conidia are spread by wind to nearby maize plants during flowering as it is known from other soil- and plant residue-borne pathogens like *Fusarium* or *Aspergillus* (57–59). After harvest, infected crop residues like husk leaves can serve as a source of inoculum when incorporated into the soil. If conditions are favorable, such as adequate moisture and temperature, *Trichoderma* may survive and potentially infect subsequent maize or wheat crops planted in the same field (20).

In conclusion, our study reveals significant heterogeneity in the pathogenicity among and within *Trichoderma* species, with *T. afroharzianum* displaying the unique ability to infect maize without wounding. This capability, coupled with its saprophytic competence, underscores its potential to cause recurrent infections and significant crop damage. In contrast, species like *T. atroviride* and *T. asperellum* rely on pre-existing wounds for infection, indicating different pathogenic mechanisms within the genus. The dual role of pathogenic *Trichoderma* strains as efficient saprophytes and pathogens highlights the need for comprehensive monitoring and management strategies. These findings emphasize the importance of understanding the diverse types of pathogenicity among and within *Trichoderma* species to effectively protect crop health and yield. Moreover, *Trichoderma* may thus be an exceptional example to explore the evolution of fungal pathogens from a saprophytic to a fully parasitic life style. Finally, our findings also highlight the importance of more careful risk assessment and quality control for biocontrol products, in order to avoid releases of microorganisms which are harmful to crops.

## Materials and Methods

### Isolation of *Trichoderma* spp. from maize samples

Maize cobs and stalks were collected from symptomatic and non-symptomatic plants of silage and grain maize fields in Germany and France from 2018 to 2023. Thirty randomly chosen kernels from each cob were surface sterilized for 10 min with 0.25% silver nitrate and placed on potato dextrose agar (PDA) with 400 µg/ml streptomycin (Duchefa Biochemie, Haarlem, Netherlands) and 30 µg/ml rifampicin (AppliChem, Darmstadt, Germany). Plates were stored at 22°C under 12h/12h light-dark cycle in climate chambers. After two days, outgrown *Trichoderma*-like mycelium was transferred to PDA plates. Individual cultures were grown for two weeks at 22°C under 12h/12h light-dark cycle in growth chambers (Mytron, Heiligenstadt, Germany) and determined morphologically under a light microscope at genus level. Single conidia cultures were produced and isolates were stored on synthetic low nutrition agar (SNA) plates at 4°C.

### Isolation of *Trichoderma* spp. from soil

Sixty soil samples were collected in 2022 either from fields with previous Trichoderma ear rot infection or non-infected fields. From each field site, four soil samples of the upper 15 cm of the soil profile were collected after removal of surface plant material. The samples were stored in a cooling chamber at 5°C until use. Soil samples were passed through a 5 mm gauze to remove coarse debris and plant material. Ten g of soil were dissolved in 100 ml water and shaken for 20 min. After sedimentation, soil suspension was diluted with sterile water to 10^-2^ and 1 ml was added to sterile PDA plates containing Bengal rose (50 ppm) and antibiotics (200 ppm Streptomycin, 200 ppm Rifampicin). Plates were incubated at 25°C with 12 h light cycle for four days. After incubation, individual colonies with *Trichoderma*-like appearance were picked with a sterile loop and transferred to PDA. Individual cultures were grown for two weeks at 22°C under 12h/12h light-dark cycle in growth chambers (Mytron, Heiligenstadt, Germany), and determined morphologically as above. Single conidia cultures were produced and stored on SNA as above.

### Isolation of *Trichoderma* spp. from biocontrol products

Samples of commercial biocontrol products were dissolved in sterile water and transferred to PDA. Liquid products were spread directly on PDA plates. The morphological characteristics of the isolates were examined to ensure that they were *Trichoderma* spp. Single conidia cultures were produced as above and stored on PDA plates at 4°C.

### Cultures and strains used in this study

*Trichoderma* strains and their origin used in this study are listed in Table A1. Among the 153 strains, nine isolates were isolated from commercially available biological fungicides and soil additives. These bio-products were directly purchased from the manufacturers.

### Plant cultivation and pathogenicity assesment

Maize seeds were sown in 18 cm diameter pots filled with a mixture of potting soil, sand and compost (3/1/3). Pots were placed in the greenhouse at 25°C temperature and 14h/10h day-/night cycle. Plants were watered as needed.

Spores from single spore cultures were transferred to PDA plates containing antibiotics, and incubated at 23°C in a growth chamber. After two weeks, sterile water was added to plates and conidia were scraped off with a microscope slide. Spore concentration was measured with a Thoma Haemocytometer (Merck, Darmstadt, Germany) and adjusted to 1×10^6^ conidia per ml. The primary cobs of maize plants were inoculated seven days after silk channel emergence (BBCH 65). A needle pin was dipped in conidia suspension and engraved in the cob through the husk, hurting the kernels. Ten plants in two repetitions were inoculated and twenty plants were inoculated with sterile water, which served as control. Four weeks (28 dpi) after inoculation, husk leaves of inoculated and control cobs were removed and percent disease severity was visually estimated (0-100%) according to EPPO Guidelines PP1/285 (60).

### Morphological characterization

Morphological observations of colonies were based on isolates grown on PDA (Merck, potato-dextrose agar) after two weeks in a growth chamber at 25 °C with alternating 12 h/12 h light/dark cycles. Photos were taken 14 days after inoculation.

### DNA extraction, PCR amplification and sequencing

Strains used in this study (Supplementary table A1) were grown on PDA for 7 d at 25 °C. Approximately 50 mg of fresh mycelium scraped from the agar surface were used for genomic DNA extraction with Quiagen Plant Mini Kit (QUIAGEN GmbH, Hilden, Germany) by following the manufacturer’s protocol. Three loci, part of the nuclear rDNA *ITS* region, 1.2-kb fragment of the *translation elongation factor 1-α* (*TEF1-α*), 1.1-kb fragment of the RNA polymerase II second largest subunit B (*RPB2*) were amplified with the primer pairs *ITS1f* (61) and *ITS4* (62), *EF1-728F* (63) and *TEF1LLErev* (64), *RPB2-5F* and *RPB2-7R* (65) respectively. PCR mixtures (25 µl) contained 2.5 μl Buffer, 5 µl MgCl_2_ (5 mM), 1 μl dNTP’s (0.2 mM), 0.4 μl of each primer (1 µM), 0.125 µl of Taq polymerase (1 U), 2 μl of DNA template and 13,575 μL of sterile deionized water. PCR reactions were performed in a Biometra Thermalcycler (AnalytiK Jena, Germany) using cycling conditions as in Gu et al. (2017) (66). PCR products were checked on 1% agarose gel electrophoresis stained with ROTI^®^GelStain (Roth, Germany). Amplicons were purified with QIAquick *PCR Purification* Kit (QUIAGEN GmbH, Hilden, Germany) and sequenced with the corresponding PCR primers in both directions by Macrogen Europe (Netherlands). Raw nucleotide sequences were edited in MEGA v. 7 (67).

### Phylogenetic analyses

The *ITS* gene was sequenced for all green shades isolates from maize and soil and used for preliminary BLAST searches in NCBI GenBank (68) in order to retain only *Trichoderma* isolates. Then *TEF1-α* and *RPB2* sequences were generated for the *Trichoderma* isolates and used for phylogenetic analysis. BLAST searches in NCBI GenBank (68) and TrichoBLAST (69) were conducted for identifying closely related species. Based on sequence similarity, additional 85 sequences were retrieved from the NCBI GenBank (Supplementary table A1). Sequences arranged in Bioedit v. 7.2.6 (70) were further aligned in MAFFT v. 7 online version (71) using the iterative refinement option G-INS-i and manually optimized with MEGA v. 7. Sequences of each locus were aligned separately and sequences of *TEF1-α* and *RPB2* were combined using Sequence Matrix v. 1.8 (72). Phylogenetic trees were constructed using maximum parsimony (MP), maximum likelihood (ML) and Bayesian inference (BI) methods. MP analysis was conducted in PAUP v. 4.0b10 (73) using heuristic searches with Tree Bisection-Reconnection (TBR), maxtrees set to 10000, ten trees saved per replicate and clade stability assessed with 1,000 bootstrap replicates. ML analyses were performed using RAxML-HPC Blackbox version 8.2.8 (74) as implemented in the CIPRES Science Gateway (75) with estimated proportion of invariable sites GTRGAMMA-I model and branch support evaluated by running 1,000 bootstrap replicates. Bayesian inference carried out in MrBayes v. 3.2.7 (76) used the best substitution model for tree reconstruction estimated by both the Akaike information criterion and the Bayesian information criterion with jModelTest 2.0 (77,78). Nucleotide substitution models in the two-gene concatenated trees were HKY + I + G for TEF1-α and SYM + I + G for RPB2. Two analyses of four Markov chains were run simultaneously starting from random trees for 5 million generations and sampling every 100th generation. After discarding the first 25% of trees as burn-in phase, a consensus Bayesian tree and Bayesian posterior probabilities (BPP) were determined based on all remaining trees. A BPP above 0.95 was considered as significant value. Trees were visualized in FigTree v. 1.4.0(79). Sequences generated in this study were deposited in GenBank and accession numbers are shown in Supplementary table A1. Alignments and trees have been deposited in TreeBASE.

### Statistical analysis

Statistical analysis was conducted using STATISTICA version 13 (Statistica GmbH, Germany). Experiments were conducted in a fully randomized design with ten plants of each treatment in two replications. Differences between means of disease severity were analyzed using parametric ANOVA by 5% probability. Analysis of variance (ANOVA) was carried out by Tukey-HSD-test at 5% probability.

## Institutional Review Board Statement

Not applicable

## Acknowledgments

We would like to thank the companies, universities and research institutions that provided us with *Trichoderma* based biocontrol agents. We would like to thank Milica Mihajlovic from the Institute of Pesticides and Environmental Protection, Belgrade-Zemun, Serbia and Monica Mezzalama from AGROINNOVA - Interdepartmental Centre of the University of Torino for providing the *Trichoderma* isolates from their strain collection. We also thank Manuela Mücke and Tobias Wille for technical support of the experiments and for performing DNA sequencing, respectively.

